# Information-dependent Enrichment Analysis Reveals Time-dependent Transcriptional Regulation of the Estrogen Pathway of Toxicity

**DOI:** 10.1101/038570

**Authors:** Salil N. Pendse, Alexandra Maertens, Michael Rosenberg, Dipanwita Roy, Rick A. Fasani, Marguerite M. Vantangoli, Samantha J. Madnick, Kim Boekelheide, Albert J. Fornace, James D. Yager, Thomas Hartung, Melvin E. Andersen, Patrick D. McMullen

## Abstract

The twenty-first century vision for toxicology involves a transition away from high-dose animal studies and into *in vitro* and computational models. This movement requires mapping pathways of toxicity through an understanding of how *in vitro* systems respond to chemical perturbation. Uncovering transcription factors responsible for gene expression patterns is essential for defining pathways of toxicity, and ultimately, for determining chemical mode of action, through which a toxicant acts. Traditionally this is achieved via chromatin immunoprecipitation studies and summarized by calculating, which transcription factors are statistically associated with the up-and down-regulated genes. These lists are commonly determined via statistical or fold-change cutoffs, a procedure that is sensitive to statistical power and may not be relevant to determining transcription factor associations. To move away from an arbitrary statistical or fold-change based cutoffs, we have developed in the context of the Mapping the Human Toxome project, a novel enrichment paradigm called Information Dependent Enrichment Analysis (IDEA) to guide identification of the transcription factor network. We used the test case of endocrine disruption of MCF-7 cells activated by 17β estradiol (E2). Using this new approach, we were able to establish a time course for transcriptional and functional responses to E2. ERα and ERβ are associated with short-term transcriptional changes in response to E2. Sustained exposure leads to the recruitment of an additional ensemble of transcription factors and alteration of cell-cycle machinery. TFAP2C and SOX2 were the transcription factors most highly correlated with dose. E2F7, E2F1 and Foxm1, which are involved in cell proliferation, were enriched only at 24h. IDEA is, therefore, a novel tool to identify candidate pathways of toxicity, clearly outperforming Gene-set Enrichment Analysis but with similar results as Weighted Gene Correlation Network Analysis, which helps to identify genes not annotated to pathways.

## Introduction

Much of what we understand about the effects of toxic compounds on human health comes from decades of experiments in animal models. This knowledge currently underwrites many of the safety regulations concerning exposures to hazardous compounds in commercial, industrial, and environmental applications. The testing strategies for these in-life animal tests are expensive, time consuming, and exorbitant in the use of animals (Cooper, Lamb et al. 2006, Hartung and Rovida 2009). Differences between human biology and laboratory animals cause difficulties in definitively assessing the safety of a compound from animal studies (Hartung 2009)). Additionally, extrapolating from high-dose conditions typically required for *in vivo* animal testing to chronic exposures relevant to human safety is problematic because of nonlinear dose-response relationships at high treatment levels. Together, these facts argue for new approaches for toxicity testing based on human biology (NRC 2007, Andersen and Krewski 2009, Andersen and Krewski 2010).

The development of *in vitro* toxicity assays and computational models can presumably replace traditional in-life animal testing. High-throughput *in vitro* screening batteries designed to assess mode-of-action and hazard are currently being used to prioritize compounds for conventional in-life testing (i.e., the EPA ToxCast and NIEHS Tox21 programs). Integrating prior knowledge about biological pathways with data from screening programs yields models that are predictive of *in vivo* testing results (Thomas, Philbert et al. 2013, Rotroff, Martin et al. 2014). However, these approaches rely heavily on knowledge of the underlying pathway of toxicity (PoT) — the mechanism by which exposure to a toxicant leads to adversity (Hartung and McBride 2010; Kleensang et al. 2014). For many commercially important chemicals their PoT are poorly understood. Thus, it would be valuable to develop a system for deriving PoT *de novo*. With this goal, the Human Toxome project (Bouhifd et al. 2015) was started in order to employ-omics technologies to start a catalogue of PoT.

Short-term full genome gene expression experiments have recently been found to be predictive of the results of 2-year bioassays (Thomas, Wesselkamper et al. 2013). This observation suggests that microarray and other high-throughput experiments could help define PoT without having to rely on incomplete and possibly misleading literature (Ioannidis 2005, Hartung 2013). To demonstrate the value of this strategy, we use estrogen receptor signaling in the well-studied MCF-7 human breast cancer cell line as a model of estrogenic endocrine disruption, which underwent International validation^1^.

Exposure to exogenous estrogens has been linked to reproductive and developmental effects, and breast and uterine cancers. Estrogens act by binding to various estrogen receptors, including ERβ, GPER, and various ERα isoforms (i.e., ERa36 and ERa46). Working in concert, these receptors orchestrate estrogen-dependent processes through regulation of transcriptional programs in various tissues. However, comparison between gene expression datasets and high-throughput chromatin immunoprecipitation (ChIP) has revealed a relatively small overlap, suggesting an incomplete picture, in which gene expression is regulated largely by cis-activation through ERs. These findings indicate that while a model of activation of gene expression through ER binding at promoters may serve as a first-order approximation of PoT—there are additional aspects that need to be considered to connect the molecular initiating event (ligand binding to the receptors) to the adverse cellular outcome (here as altered proliferation).

In addition to estrogens acting directly through ERα and ERβ, there is increasing evidence for regulatory contributions from additional transcription factors (O’Lone, Frith et al. 2004). ERα interacts with a number of transcriptional modulators, including AP-1 (Zhao, Gao et al. 2010), Sp1 (Schultz, Petz et al. 2003), SNCG (Jiang, Liu et al. 2003), and Sin3A (Ellison-Zelski, Solodin et al. 2009). Non-genomic signaling, originating from estrogens binding to the G-protein-coupled receptor GPER or from ERα isoforms anchored to the plasma membrane, initiates kinase cascades that drive transcription through heretofore-unknown mechanisms.

Predicting the transcription factors responsible for a cellular response would significantly contribute to PoT identification (Essaghir, Toffalini et al. 2010, Maertens, Luechtefeld et al. 2015). However, traditional approaches for identifying transcription factors from gene expression patterns use data from a small subset of the genome. Here, we investigate the transcription factor network responsible for estrogen-mediated transcriptional changes using a novel approach that makes use of a higher proportion of the biological information than conventional methods. We have performed gene expression microarray experiments exposing the MCF7 breast carcinoma cell line to the canonical estrogen, 17β-estradiol (E2). By combining the observed gene expression changes with publically available ChIP data, we generated a putative gene-regulatory network.

## Methods

### Cell culture

MCF-7 cells were seeded at a density of 300,000 cells/well in 6-well plates and allowed to grow for 72 hours in complete growth media composed of DMEM/F12 media supplemented with 10% fetal bovine serum (FBS, Atlanta Biologicals, Flowery Branch, GA), non-essential amino acids, 10μg/mL bovine insulin and gentamicin. After 72 hours, cells were rinsed with PBS and placed in treatment media composed of DMEM/F12 supplemented with 5% dextran charcoal-stripped fetal bovine serum (DCC, Gemini Bio-products, Sacramento, CA, US, no. 100-119), nonessential amino acids, 6ng/mL bovine insulin and gentamicin for 48 hours. Cells were then exposed to 17β estradiol (E2, SigmaAldrich, St. Louis, MO, USA, no. E8875) or vehicle control dimethylsulfoxide (DMSO, Sigma Aldrich, no. D8418) in fresh treatment media for 2, 4, 8, and 24 hours. Samples were scraped into TRI Reagent (Sigma Aldrich, no. T9424) and stored at-80°C until RNA isolation and q-PCR analysis.

### Gene expression microarray experiments

Total RNA from MCF-7 cells was extracted using TRizol Reagent according to manufacturer’s instruction, and purified using RNeasy Mini Kit (Qiagen). Purified RNA was quantified by using NanoDrop ND-1000 spectrophotometer and the quality of RNA was analyzed by using Agilent Bioanalyzer (Agilent). 100 ng of total RNA from treated and control cells were converted into cDNA and then into labeled cRNA using Agilent LowInput QuickAmp Labeling Kit (Agilent). The resulting cRNA was labeled with Cy3. Labeled cRNAs were then purified, and RNA concentration and dye incorporation were measured using NanoDrop ND-1000 spectrophotometer. Hybridization to Agilent SurePrint G3 human whole genome 8x60K microarray (Agilent) was conducted following manufacturer’s protocol. Microarrays were scanned with an Agilent DNA microarray scanner. Feature Extraction (11.5.1.1 version, Agilent) was used to filter, normalize, and calculate the signal intensity and ratios. Processed data were subjected to GeneSpring (Agilent) analysis.

### Gene expression analysis

Data from microarray experiments was analyzed using GeneSpring (Agilent) software. Raw data were imported and quantile-normalized. Fold-change expressions for the probes were calculated by calculating the ratio of change from probes in time-matched controls. Significance for the change was computed using a t-test and corrected for multiple tests using FDR correction. Genes were then assigned to their respective probes using the annotation files created by Agilent for the microarray plates used.

### Transcription factor database curation

Several compendia exist of transcription factor-target interaction that we can use to uncover a regulatory network, through which estrogen acts. For this study, we used Chip-X Enrichment Analysis (ChEA) database (Kou, Chen et al. 2013) and the Encode database (Bernstein, Birney et al. 2012). The databases were combined to increase coverage of the databases. This combined database was used to calculate enrichment using IDEA and Gene-Set Enrichment Analysis (GSEA).

### Information Dependent Enrichment Analysis

The workflow for calculating enrichment using IDEA is represented in **Figure 1 Error! Reference source not found..** Consider the set of *N* genes identified as upregulated. Let *δ_i_* describe the relationship between a transcription factor and gene *i,* where *0≤i≤N* and *δ_i_* = *1* if the transcription factor regulates gene *i*. From this group, a set {*E_n_*} of the *n* most highly up-regulated genes can be defined. This sample of *n* genes contains *Σ_{i<n}_ δ_i_* genes regulated by the transcription factor. Fisher’s exact test provides the probability *f_n_* that a gene selected from {*E_n_*} and a gene selected from the genome at large have equivalent likelihood of being regulated by the transcription factor. *f_n_* can be calculated for all values of *n* and test statistic *t* defined as the min*_n_ fn*.

**Figure 1.**
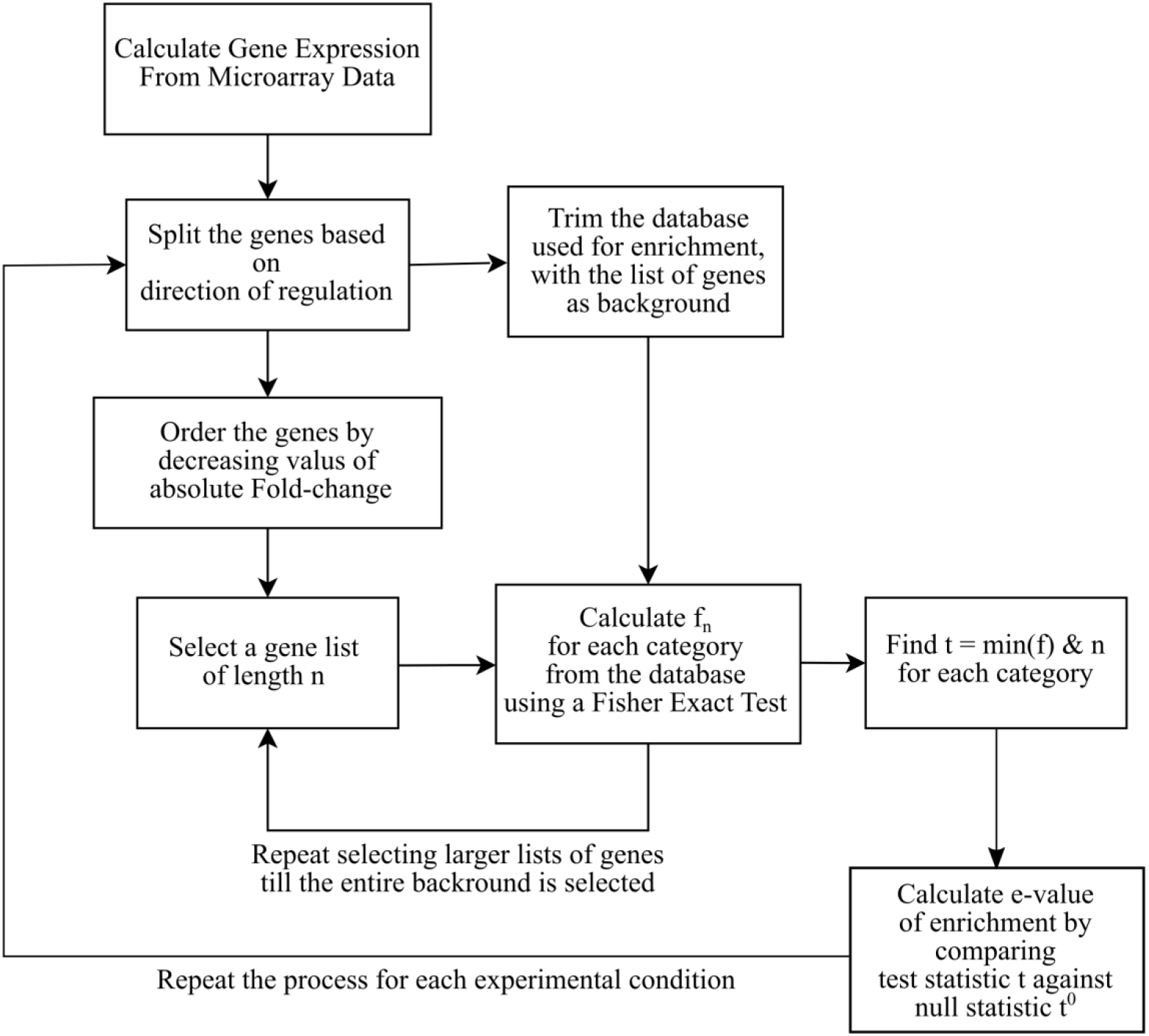
Flowchart for computing enrichment using the IDEA Algorithm

**Figure 2.**
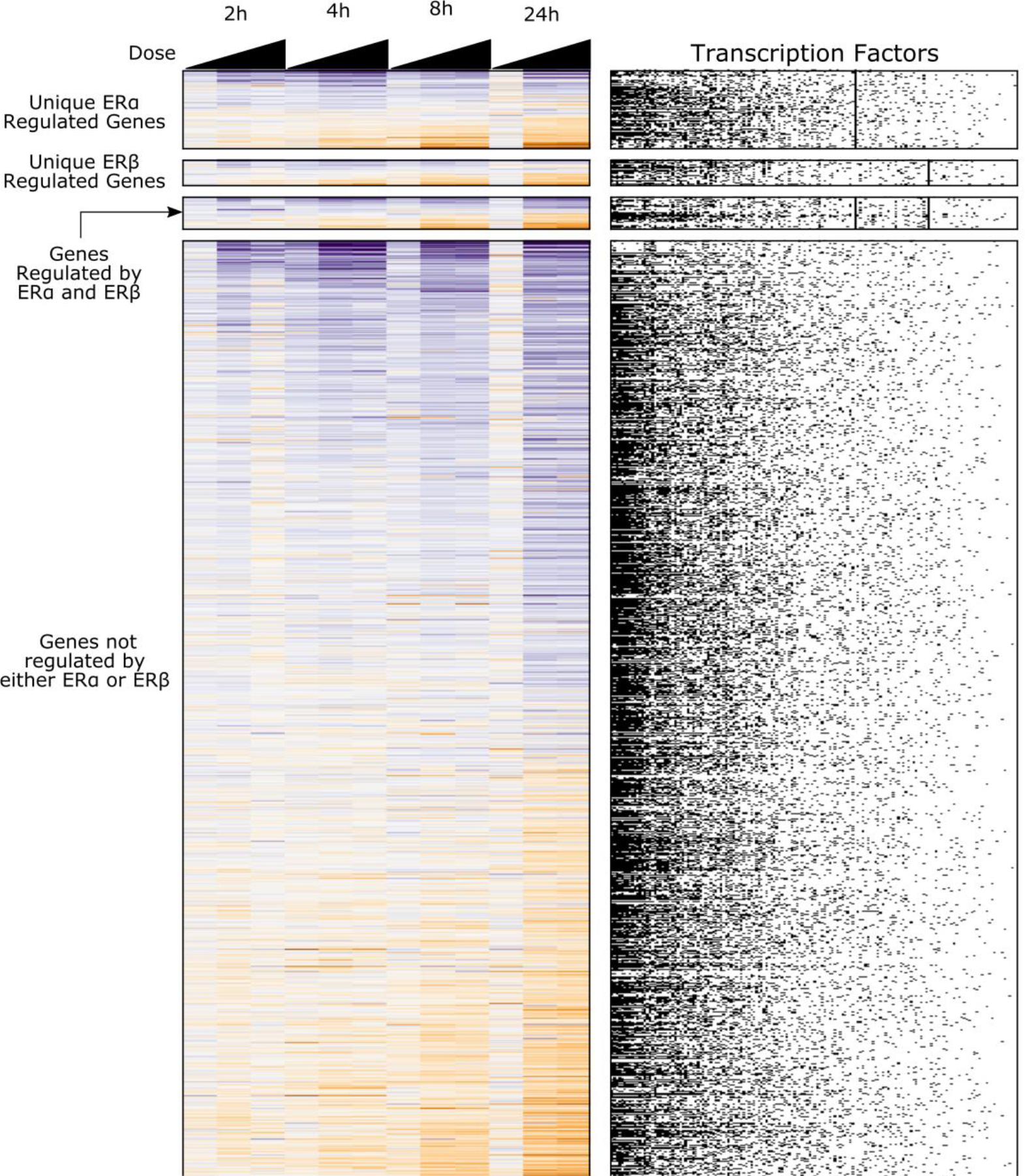
Genomic Response of E2-treated MCF7 cells. (A) Matrix of expression changes for all differentially expressed genes. Each row indicates a gene, each column indicates a treatment condition increasing in dose from left to right. A small fraction of the differentially expressed genes have been shown to bind either ERα or ERβ in previous studies. (B) Transcription Factor Regulation Matrix. Each row is a gene and each column is a transcription factor. Black dots indicate that gene is shown to have the corresponding transcription factor binding.

To determine whether the transcription factor is associated with up-regulated genes, we used Monte Carlo hypothesis testing. A distribution {*t^0^*} of null-model test statistics was established by permuting the N upregulated genes and calculating *f_n_^0^* for each permutation (Supplemental Figure S1). The best estimate of the probability *e* that *t* is consistent with the null model is determined by quantile function of {*t^0^*} (Supplemental Figure S2).

This procedure was repeated for all transcription factors in the database and for both up-and down-regulated gene sets. The Benjamini-Hochberg multiple test correction was applied to the *e*values. The same procedure was repeated with KEGG and Reactome ontologies establish effects of estrogen exposure on biological processes.

### Weighted Gene Correlation Network Analysis (WGCNA)

A signed weighted gene correlation network analysis (WGCNA) network (Langfelder and Horvath 2008) was generated on the 7000 most highly expressed genes at 8h as determined by rank means expression. The network was derived based on a signed Spearman correlation using a β of 8, and clustered into modules using dynamic tree cut with a height of 0.25 and a deep split level of 3, and a reassign threshold of 2. The eigenmodules - essentially the first principal component of the modules - were then correlated with dose. Each module that had a statistically significant correlation with dose was analyzed for transcription factors using the ChEA 2015 dataset accessed via EnrichR (Chen Y, et al 2013) restricted to MCF-7 cells.

## Results and Discussion

### Response of MCF-7 cells to estrogen

The gene expression response of the MCF-7 breast carcinoma cell line to 17β-estradiol (E2) has been extensively documented (O’Lone, Frith et al. 2004). However, the bevy of gene expression studies in the MCF-7 experimental system (Rae, Johnson et al. 2005, Carroll, Meyer et al. 2006, Chang, Frasor et al. 2006, Creighton, Cordero et al. 2006, Fan, Yan et al. 2006, Frasor, Chang et al. 2006, Gaube, Wolfl et al. 2007, Kininis, Chen et al. 2007, Lin, Vega et al. 2007, Lin, Reierstad et al. 2007, Bourdeau, Deschenes et al. 2008, Chang, Charn et al. 2008) display a large degree of inconsistency, when analyzed at the gene level (Ochsner, Steffen et al. 2009, Jagannathan and Robinson-Rechavi 2011). Here, we build on this literature by performing gene expression microarray analysis on MCF-7 cells with 0.01, 0.1, and 1nM E2 for 2, 4, 8, and 24h.

The number of genes identified as differentially expressed (FDR-corrected p-values less than 0.05) in cells treated with 1nM E2 varied substantially with time and concentration—from zero genes after 2h exposure to 4113 genes after 24h **(Error! Reference source not found**.,Table 1). Interestingly, this increase in number of differentially expressed genes is not monotonic, with 547 genes identified at 4h and only 4 genes identified at 8h post treatment. Because the identification of differentially expressed genes depends on experimental factors that drive statistical power, it is unclear whether this decrease in differentially expressed genes with time is biologically meaningful. This observation—possibly an artifact—is common to functional genomics experiments that test thousands of hypotheses in parallel and here it poses a challenge for correctly interpreting data.

**Table 1.** Number of genes differentially expressed following E2 treatment in MCF7 cells. The number of genes differentially expressed was not a linear function of either time or dose.

|  | 2h |  | 4h |  | 8h |  | 24h |  |
| --- | --- | --- | --- | --- | --- | --- | --- | --- |
|  | Up | Down | Up | Down | Up | Down | Up | Down |
| 0.01 nM | 3 | 9 | 48 | 39 | 14 | 14 | 0 | 0 |
| 0.1 nM | 205 | 120 | 670 | 312 | 229 | 180 | 1350 | 954 |
| 1.0 nM | 0 | 0 | 375 | 172 | 3 | 1 | 2101 | 2012 |

Because the lists of differentially expressed genes derived from microarray experiments are determined by statistical power (e.g., number of replicates, RNA isolation protocol, microarray platform, etc.) as well by biology, using standard over-representation analysis in this case would likely be unsound. Classical overrepresentation analysis—commonly used to assign functional ontology descriptions to sets of genes—suffers from a number of shortcomings. Gene expression changes are aggregated into lists of up-and downregulated genes based on significance criteria, magnitude of change, or some combination of these and other factors. These choices result in a list of the most extremely responding genes. Rarely do genes encode a protein that is solely responsible for the cellular response to stimuli. As such almost always multiple genes are transcribed to varying degrees to bring about a response and using some arbitrary cut-off does not take into account this nature of gene transcription. More critically for toxicology, this methodology would likely miss subtle effects at low doses or early time-points, when very few genes are identified as differentially expressed. This approach often results in no enrichment observed at low exposures and trivial categories enriched at high exposures. The number of genes used for calculating enrichment is dependent completely on how many are identified as significant. This makes it impossible to compare relative enrichments between categories as inherently different lists were used to perform the computation.

To address this problem, we developed a novel approach for assessing enrichment from high-throughput data, which has the advantage of being relatively insensitive to variability in statistical power in assignment of differential expression and makes use of a larger complement of the gene expression data to determine enrichment for transcription factor binding or functional ontology.

### The IDEA algorithm for enrichment analysis

Our novel approach bypasses the pitfalls of existing methods by avoiding differentially expressed gene lists and instead using the entire set of microarray data to create a gene list ordered by expression values. This avoids attempts to balance the sensitivity and specificity with statistical cut-offs and concerns about bias in the background.

Slicing gene lists of different length allows us to look at how the strength of enrichment for each category changes with increasing number of genes. If a transcriptional network is active, then even at low exposures multiple genes belonging to that network will be expressed, albeit not strongly. Furthermore, those genes will follow a pattern of expression that will be captured by their ordering **(Figure 3 Error! Reference source not found.).** This will give the statistical tests used enough power to test for enrichment when a large gene list is selected. Each enrichment for category is computed over the same background and using the same number of genes. This approach allows us to easily compare the relative contribution of categories to overall signal by looking at number of genes required to achieve peak enrichment. With this information, and how the relative ordering of genes required changes across experimental conditions, we can predict how the cell responds to stimuli in a time and dose dependent manner. That allows us to discern the signaling cascade of transcription factors and its evolution over increasing exposure. In order to better understand the changes in transcription factor enrichment, we also created a visualization tool to interact with the IDEA results(Link to visualization on the web).

**Figure 3.**
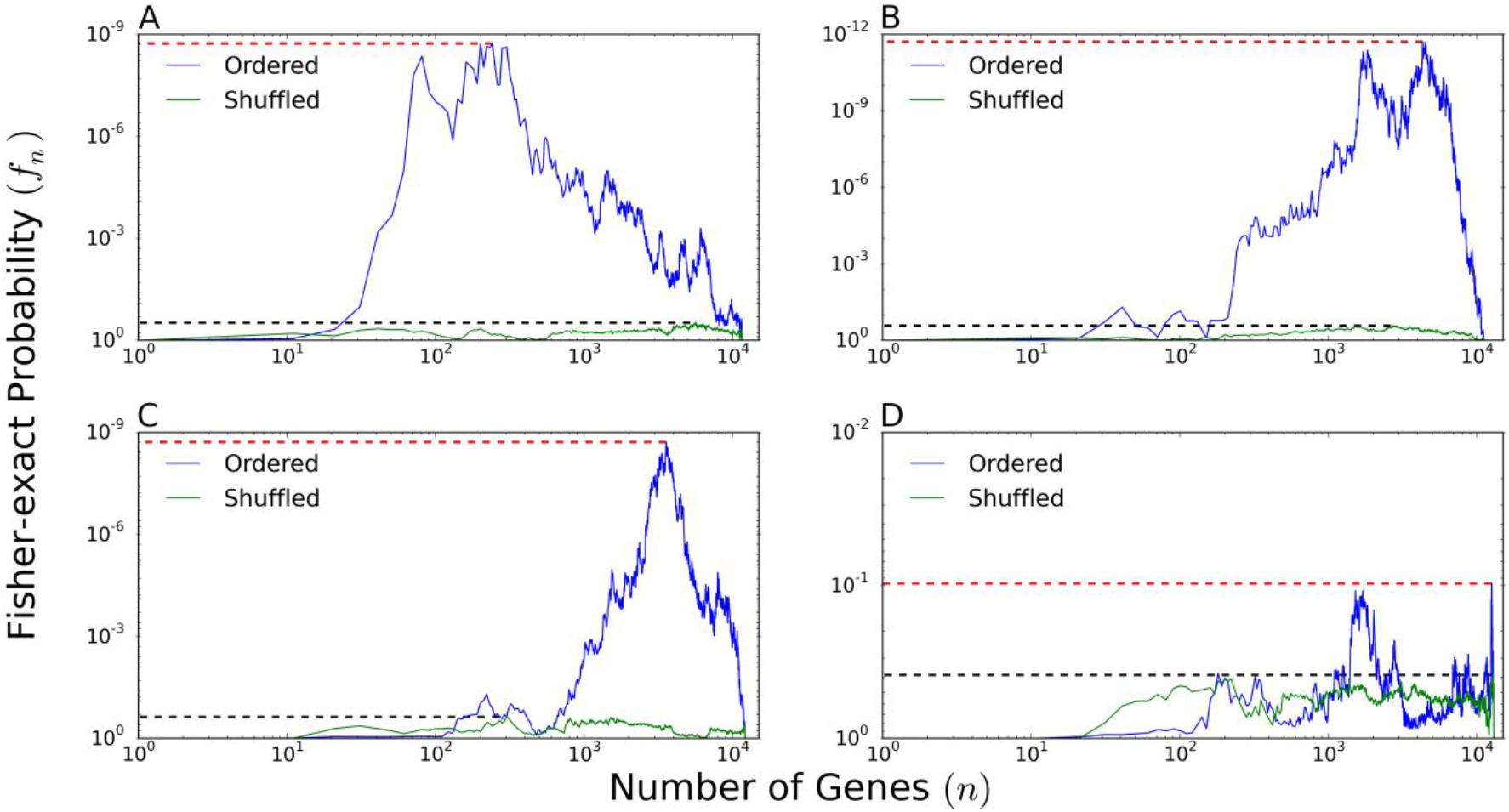
Plot of p-values determined by Fisher Exact test for genes regulated by ERα in MCF7 cells. A) Genes up-regulated 24h post-treatment, (B) downregulated 24h post-treatment, (C) up-regulated 2h post-treatment and (D) down-regulated 2h post-treatment

### Estrogen receptors drive the short-term transcriptional response to E2

We surveyed existing ChIP datasets to better understand the factors driving estrogen-mediated transcriptional changes. Interestingly, ERα and ERβ—two primary estrogen transcription factors—have only been shown to bind a small fraction of differentially expressed genes **(Error! Reference source not found.).** This low degree of overlap between receptor binding and gene expression regulation is in sharp contrast to other nuclear receptors that have been shown to account for approximately half of the affected genes’ expression (van der Meer, Degenhardt et al. 2010, McMullen, Bhattacharya et al. 2014).

At 2h, of all the up-regulated genes, 28% are needed for peak ERα and ERβ enrichment. At 24h this ratio shifts to only 1.7 % for ERα and 7.8% for ERβ. In other words, as would be expected, the ERα and ERβ signal strengthens over time - at 2 hours it is barely over background level while at 24 hours the signal is much more distinct. Most importantly, from the point of view of the bioinformatician looking to discern a signature of toxicity, this approach allows quantifying the strength of the signal at any given time point.

We investigated the large variation in number of genes needed for peak enrichment to determine whether this factor contained information about the biology of the system. Similar sets of ERα and ERβ genes are up-regulated at 2h and 24h (Figure 4). Also the highest responding genes at 24h are also up-regulated at 2h. Hence the difference in enrichment comes from change in expression patterns of similar genes at 2h and 24h.

**Figure 4.**
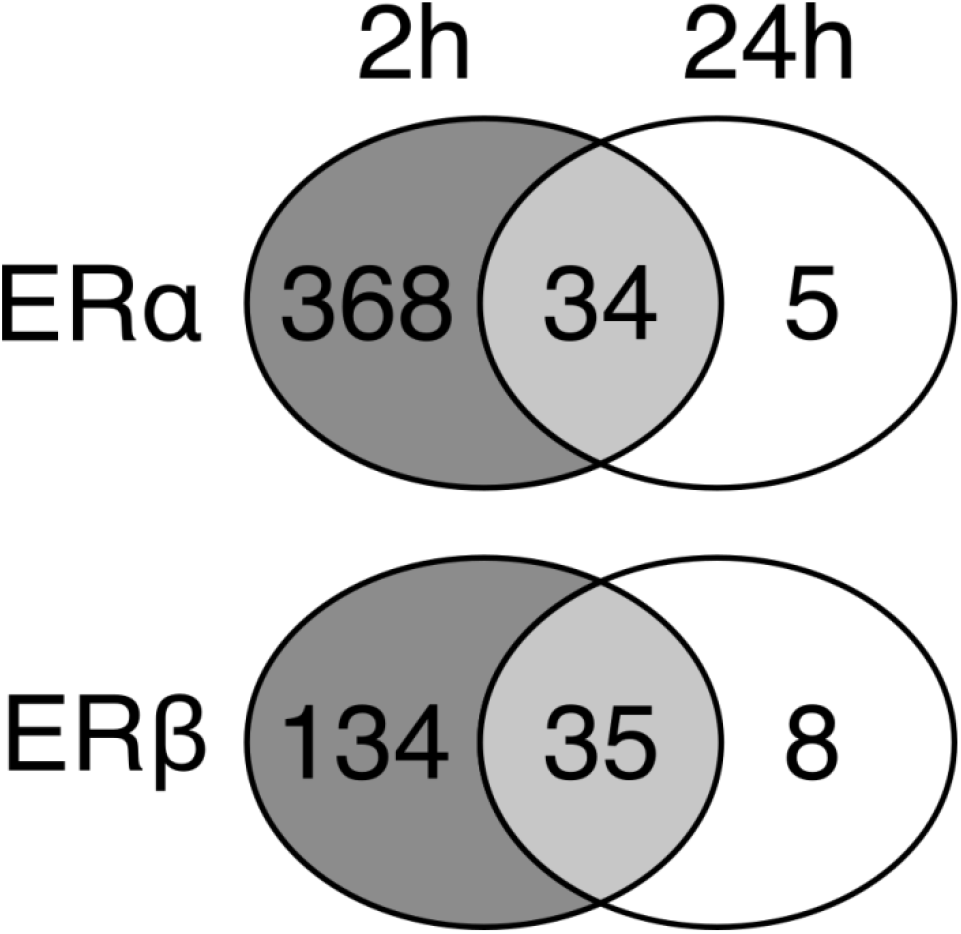
Overlap between genes enriched and bound by ERα and ERβ at 2h and 24h. Almost all genes contributing to enrichment at 24h were also expressed at 2h.

Plotting the distribution of expression for all ERα and ERβ regulated genes (Figure 5) reveals that at 24h, there is a larger set of highly up-regulated genes (greater than 4-fold change) than in background. Alternatively at 2h, all the ERα and ERβ regulated genes follow background pattern of gene expression more closely. This result indicates that at ERα and ERβ are strong drivers of the underlying expression pattern at 24h but their signal is mediated only by a limited subset of the full group of differentially expressed genes.

**Figure 5.**
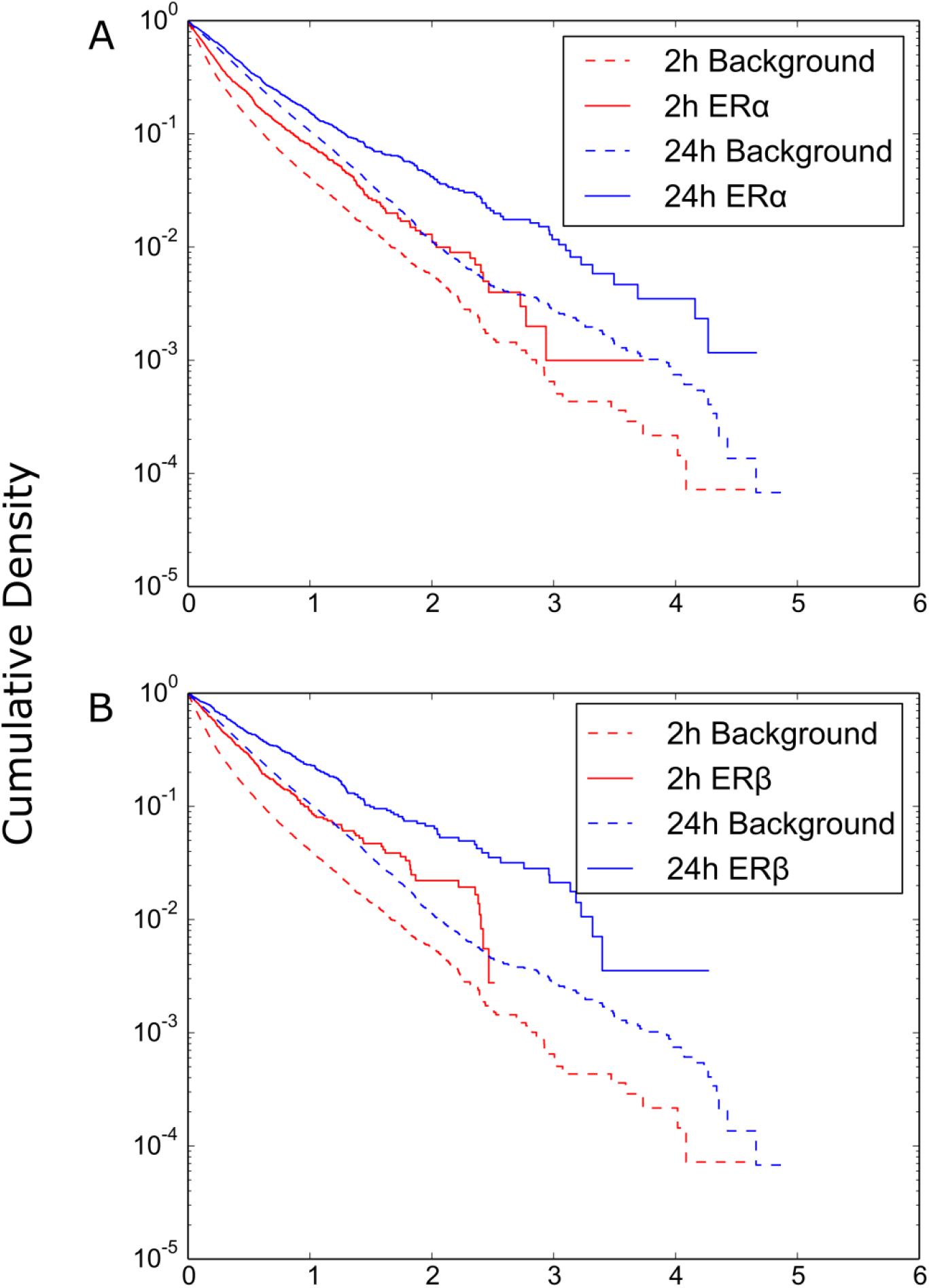
Cumulative distribution of expression changes in upregulated genes that are regulated by (A) ERα and (B) ERβ at 2h (red) and 24 h (blue). Background distributions (dashed) reflect all upregulated genes at the given condition. More ERα and ERβ bound genes have greater than 2 log2 fold change (solid blue line) at 24h than any other genes at any other condition

A very different pattern of enrichment in observed in the case of down-regulated genes. Looking at gene expression at 2h, we see no enrichment in ERα and ERβ profiles. We observed enrichment for a few other transcription factors, like ZNF217, indicating that the absence of enrichment is due to lack of ER signaling in these genes rather than a lack of gene expression. At later times, estrogen receptors are highly enriched by down-regulated genes. However, their peak enrichment never requires less than 37% of all down-regulated genes.

### Cell cycle alterations at 24h

Persistent exposure of MCF7 cells to E2 induces cell proliferation. Time course of estrogen receptor enrichment shows a shift from a large suite of ER genes expressed at low level to a small set of highly expressed ER genes. We believe this to be a result of a shift from estrogen-specific signaling to generic cell cycle signaling driven by ER genes. Additional evidence for this process is provided by the enrichment of E2F7, E2F1 and Foxm1 only at 24h. Both Foxm1 and E2F1 are transcriptional activators involved in cell proliferation (Stender, Frasor et al. 2007, Real, Meo-Evoli et al. 2011). E2F7 on the other hand represses the activity of E2F1 by binding to E2F1-responsive genes (De Bruin, Maiti et al. 2003, Liu, Shats et al. 2013). However at 24h, E2F7 is only enriched in the set of up-regulated genes providing further evidence to the generic cell cycle signature at later times.

In contrast, ZNF217, a transcription factor that has been implicated in cell division and differentiation in many cancers (Zhu, Zhu et al. 2005, Littlepage, Adler et al. 2012, Rahman, Nakayama et al. 2012), is enriched at all time-points among down-regulated genes. High levels of ZNF217 mRNA is a marker of poor prognosis in breast cancer (Littlepage, Adler et al. 2012). ZNF217 primarily acts by repressing genes that halt cell cycle, thereby promoting cell growth and differentiation (Thollet, Vendrell et al. 2010). This loss of gene expression is consistent with the role of ZNF217 as a repressor that is essential to proliferation in breast cancer cells (Thollet, Vendrell et al. 2010).

### Relationship to Gene Set Enrichment Analysis

Gene-set Enrichment Analysis (Subramanian, Subramanian et al. 2005) is a well-known algorithm that attempts to use a priori gene set information to calculate enrichment of gene lists. We applied the widely-used GSEA algorithm to our combined transcription factor database to better understand the relationship between IDEA and existing methods (Subramanian, Subramanian et al. 2005). The results agree closely with those obtained by us (supplementary table). At 2h, only 3 transcription factors are enriched in cells treated with 1nM E2 using recommended parameters at a FDR of less than 25%, the value used by the creators of GSEA as a valid cut-off for establishing enrichment (Subramanian, Subramanian et al. 2005). At 24h, 69 transcription factors are identified as enriched in treated cells. Additionally at 2h, ERα and ERβ are highly enriched for treated cells, whereas at 24h the E2F family of proteins is highly enriched for the treated cells. This is in line with our hypothesis that there exists a proximate ER network that then feeds into the generic cell cycle processes to effect proliferation and other phenotypic alterations associated with E2 treatment.

GSEA is geared towards discerning differences in enrichment between two experimental conditions (in toxicology studies, often a treatment and a control) by attributing enrichment of each gene set to one of the two conditions. When using GSEA to compare enrichment between Estrogen-treated and untreated cells, transcription factors associated with down-regulated genes and those that have no effect are both identified as enriched in untreated cells. As shown above, ERα and ERβ are associated with large sets of both upregulated and downregulated genes. The similarities and differences in the composition of these gene sets and their expression patterns are essential in uncovering the underlying transcription factor network. At longer exposures, GSEA identifies enrichment in ERα and ERβ in untreated cells but ignores the small set of highly upregulated genes driven by these transcription factors. This rigidity inherent to the GSEA method hinders its utility in interpreting results whereas the same transcription factors may be responsible for both activation and repression of genes through different pathways. Finally, the results of GSEA are dependent on the choice of weight function for calculating the enrichment statistic, whereas IDEA relies on the statistics of the hypergeometric distribution to calculate enrichment.

### Evidence for cell cycle signaling from functional ontologies

Cell cycle is controlled by a large number of transcription factors. Hence it was necessary to ensure that enrichment in E2F family of proteins is indicative of global cell cycle signaling in the cell. Functional ontologies like Kyoto Encyclopedia of Genes and Genomes (KEGG) and Reactome attempt to assign genes to functional categories based on information curated from experimental results. These databases are better at identifying processes (i.e., cell cycle, metabolism, etc.) that depend on a relatively large section of the genome to be expressed. As such they are an ideal complement to transcription factor databases that capture processes regulated by a small subset of genes in the genome.

To investigate the hypothesis of altered cell cycle signal appearing only after longer exposures, we calculated the enrichment of functional categories in both Reactome and KEGG using the IDEA algorithm. We observed a very clear temporal pattern of enrichment of cell cycle-related categories. At 2h and 4h, none of the cell cycle-related categories were enriched. However at 8h, we observed enrichment of some cell cycle-related categories like DNA replication. Finally at 24h post treatment, all mitotic cell cycle-related categories in both Reactome and KEGG were significantly enriched (link to visualization). Furthermore, the numbers of genes needed for peak enrichment at 24h were less than those needed at 8h, indicating stronger information content in the enrichment signal at 24h. We also clustered the enrichment profiles obtained from KEGG ontology using hierarchical clustering algorithm with Euclidian distance metric between decimal logarithms of t-values (Figure 6). The clustering showed a similar response with signaling pathways being activated as early as 2h and 4h. Cell Cycle and DNA replication were only enriched at 8h and 24h. Figure 7 illustrates the time-dependence of key transitions in transcription factor and functional ontology enrichment patterns.

**Figure 6.**
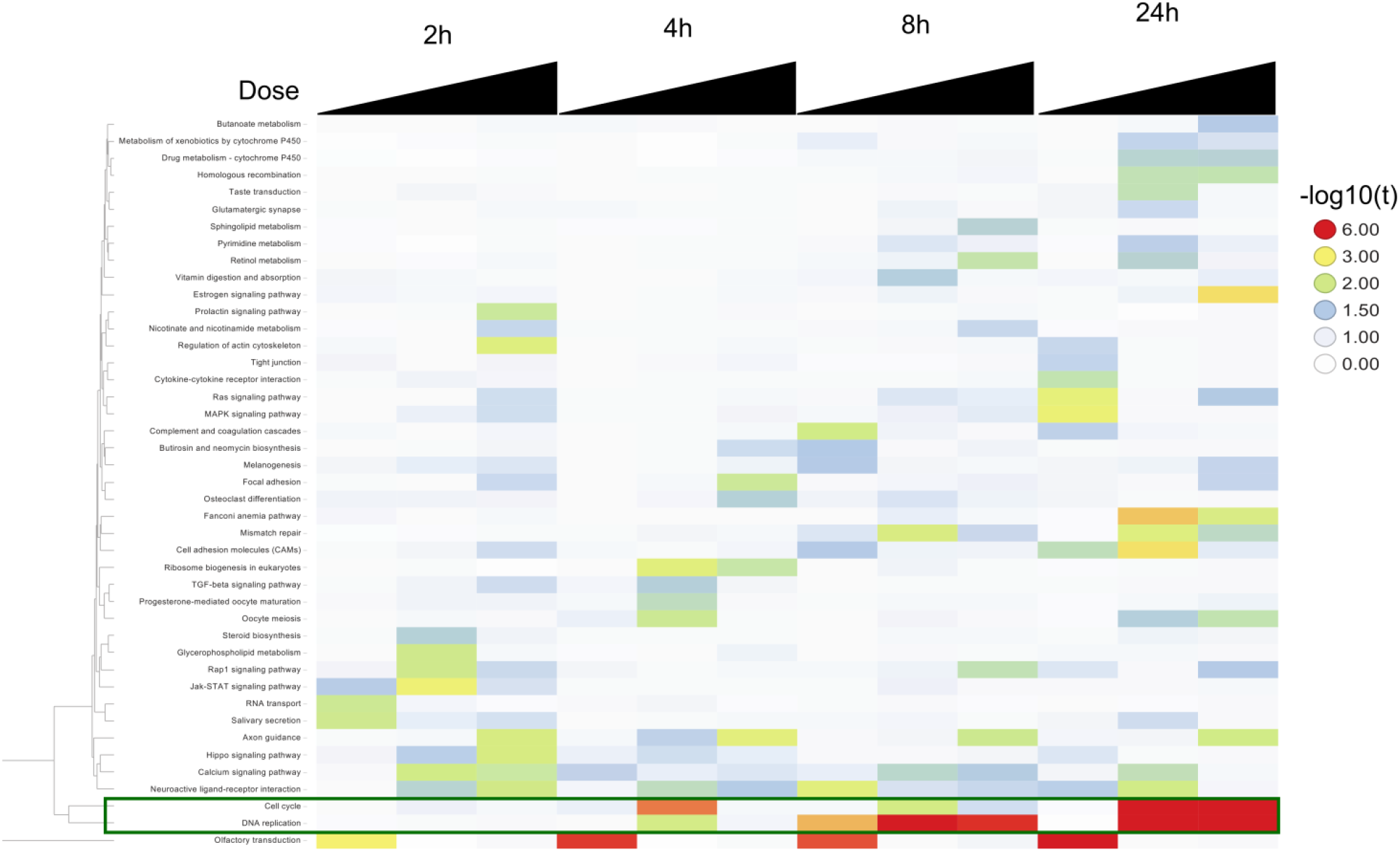
Clustering t-values for KEGG pathways. Cell cycle-related pathways cluster independent of all other pathways (shown in a green box). Cell cycle is strongly enriched only at 24h post treatment, while DNA replication is enriched at both 8h and 24h post treatment.

**Figure 7.**
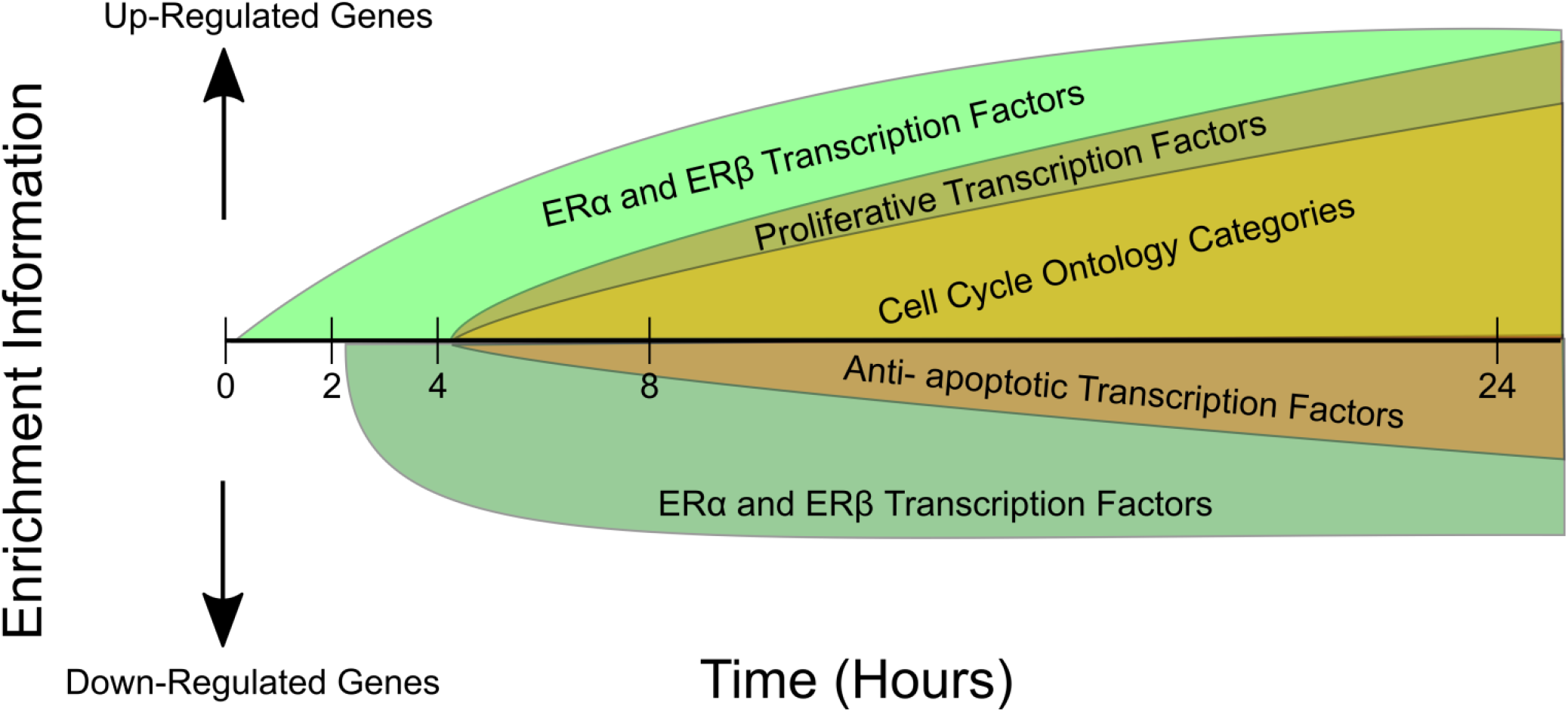
Key events in estrogen signaling. Genes up-regulated by ERα and ERβ already show a strong expression pattern at 2h post-treatment. This pattern continues to get stronger with time. Generic proliferative transcription factors like E2F1 and E2F4 are enriched for the first time only at 8h post exposure. At the same time, genes involved in cell cycle and proliferation also show a strong expression pattern. Some ERα and ERβ genes are down-regulated in response to E2 treatment, but their expression patterns do not evolve over time.

Gene expression changes are aggregated into lists of up-and down-regulated genes based on significance criteria, magnitude of change, or some combination of these and other factors. These choices result in a list of the most extremely responding genes, dependent completely on how many are identified as significant.

### *De novo* network analysis of estrogen perturbation

To investigate the data from a methodology that is blind to *a priori* knowledge of transcription factor binding sites and is relatively insensitive to concerns about technical bias, we used weighted gene correlation network analysis (WGCNA) to build a *de novo* network from the data using the dose response curve at eight hours – notably a time point where only four genes were significantly expressed in response to E2. Correlation methods offer an additional alternative to using differentially expressed genes for downstream analysis, as they take advantage of a larger portion of the data and allow for the investigation of links between genes.(Maertens et al 2015). Moreover, WGCNA assigns genes to modules based on a graph theoretical algorithm and tests for significance between the modules and experimental factors (here, the dose-response curve). The added value for identifying PoT has been recognized earlier (Andersen et al. 2015, Rahnenführer and Leist 2015).

Despite a relatively weak signal in terms of differentially expressed genes at that the 8-hour time-point, the network derived from the data contained 5 modules that were significantly correlated with E2 concentration. To understand the relationship between the transcriptional factors co-ordinating the gene expression and the modules, each module was analyzed for transcription factors using the ChEA dataset but restricted to MCF7 cells. In addition to ESR1 and ESR2, there were also well-known estrogen response pathway transcription factors such as GATA3 and cancer-related transcription factor HIF1. Moreover, both ZNF217 and TFAP2C were identified as a significant transcription factor in each module correlated with dose and as expected several transcriptional modules coincident with the enriched transcription factors for upregulated genes identified with IDEA (Supplementary Table 4).

While TFAP2C is not annotated to estrogen-responsive pathways in either the KEGG estrogen signaling pathway (Kanehisa and Goto 2000) nor does it have any GO Annotations relating to estrogen, it is a key regulator of hormone responsiveness at multiple levels. It acts both directly by regulating ERα transcription and indirectly by recruiting key estrogen pioneer transcription factors GATA3 and FOXA1, and additionally by modulating several downstream signaling pathways (Cyr, Kulak et al. 2015). *In vitro,* TFAP2C attenuation leads to a lack of mitogenic response to estrogen and *in vivo* decreased hormone-responsive tumor growth of breast cancer xenografts (Woodfield, Horan et al. 2007). Clinically, higher TFAP2C scores correlates with poorer survival for breast cancer patients (Perkins, Bales et al. 2015). Moreover, both TFAP2C and ZNF217 gene expression levels were correlated with estrogen receptor status in breast cancer dataset from TCGA, indicating that the significance of these genes for *in vivo* biology.

SOX2 was also identified in several of the modules. One key step in the regulation of breast tumor-initiating cells takes place when ERα down-regulates miR-140 (Zhang, Eades et al. 2012), which in turn increases levels of SOX2. SOX2 is considered a key regulator of stem-cell self-renewal and specifically in breast cancer tissue is thought to promote non-genomic estrogen signaling while simultaneously acting to amplify estrogen’s signal by increasing the nuclear levels of phospho-Ser118-ERα (Vazquez-Martin, Cufi et al. 2013). Expression of SOX2 is increased in early stage breast tumors, and over-expression of SOX2 increased mammosphere formation, while SOX2 knockdown prevented mammosphere formation and delayed tumor formation in a xenograft tumor initiation model (Leis, Eguiara et al. 2012). Both TFAP2C and SOX2 were also enriched using the IDEA algorithm.

The concordance between the IDEA algorithm (which also identified TFAP2C, ZNF217, and SOX2) and the transcription factors identified by WGCNA shows that both the methods complement each other and further investigation of expression analysis using WGCNA would help identify estrogen-responsive genes not annotated to ER pathways.

## Conclusions

The technologies driving modern biology produce a surfeit of data, often spanning the breadth of the genome. However, methods for extracting biological insight from the results of these experiments have lagged behind. New computational tools and visualization strategies are required to fully realize the potential of systems biology for revolutionizing toxicity testing and mapping toxicity pathways. High-throughput tools often implicate large lists of genes for particular phenotypic responses. However, translating this information into biological knowledge remains a fundamental challenge. There also exists a persistent perception in modern biological research that more information automatically leads to more knowledge. However, the quantity and complexity of high-throughput data is typically not directly translatable into advances in understanding.

Summarizing changes to transcriptional programs by associating them with existing literature and curated databases is a key modality for summarizing and understanding the results of high-throughput experiments. Here, we treated MCF7 cells with E2 and calculated transcription factors over-represented by expressed genes. We also inferred the functional implications of those gene expression changes. Because existing enrichment approaches were insufficient for interpreting these changes, we derived a novel technique that makes more complete use of the biological data.

Our tool, IDEA, provides a framework for observing patterns with gene expression studies. Toxicants with similar modes of action are expected to induce similar patterns of transcriptional change. However, changes in individual genes are typically not as robust as changes at the pathway level. Because it summarizes gene expression changes into a small subset of transcription factors or ontology categories associated with the up-and/or downregulated genes, this is a promising tool for identifying mode of action.

By considering the time-course of genes regulated by various transcription factors, we hypothesize that response to estrogen involves two distinct steps (Figure 8). During the first stage, at 2h to 4h post-treatment, signaling is dominated by cis-regulation of ERα and ERβ. This primes the cells for growth. At longer exposures, only a subset of ERα-and ERβ-controlled genes is highly up-regulated. Simultaneously, a large set of genes regulated by cell signaling transcription factors, including E2F1, E2F4, and Foxm1 are upregulated. At longer exposures, cell cycle-related categories in KEGG (Kanehisa and Goto 2000) and Reactome (Fabregat, Sidiropoulos et al. 2015) are enriched in up-regulated genes, while apoptotic and anti-proliferative categories are enriched in down-regulated genes.

**Figure 8.**
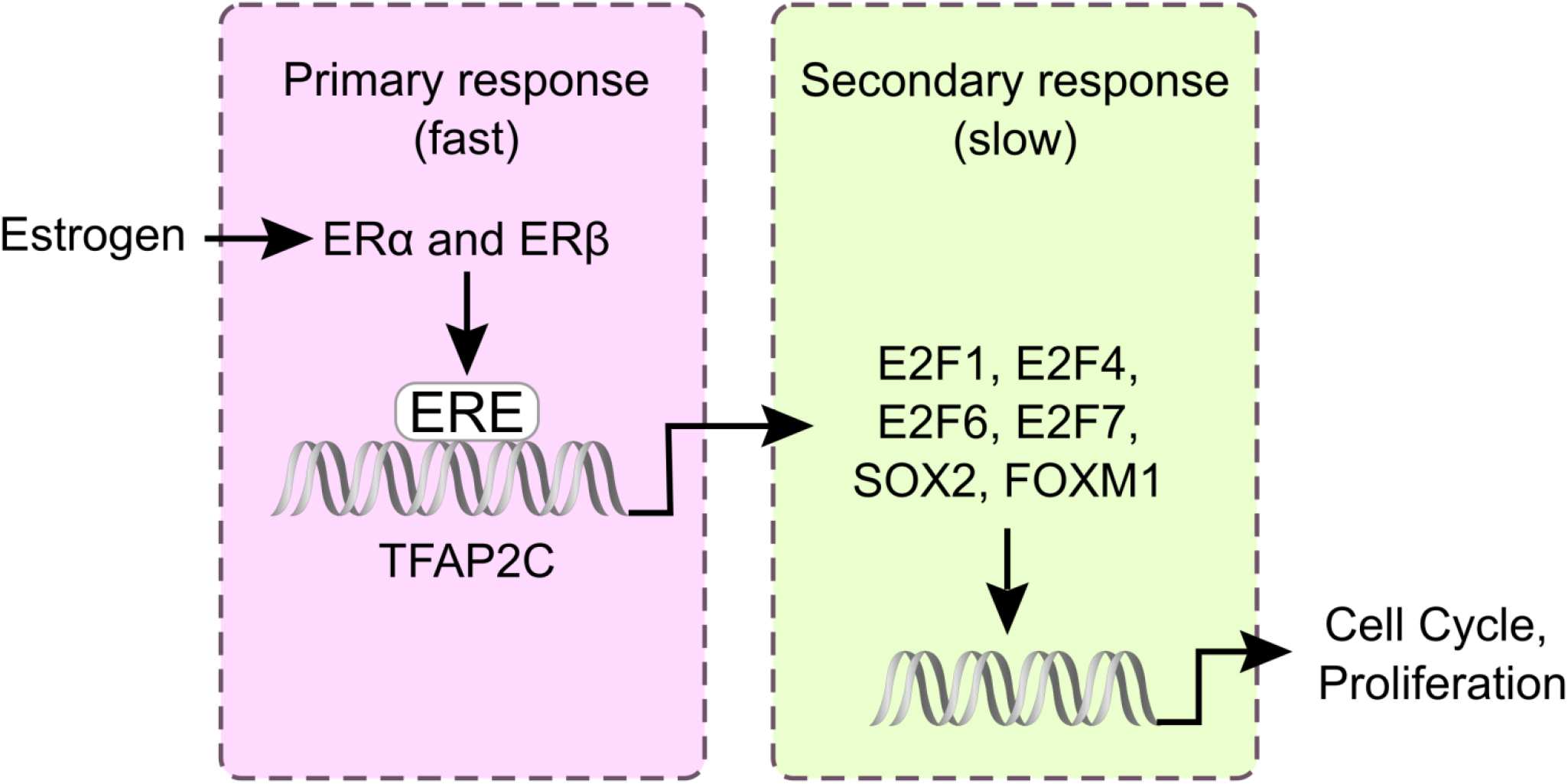
Putative transcription factor network for Estrogen signaling. ERα and ERβ bind directly to DNA via Estrogen Responsive Elements (ERE). This initiates transcription of a set of estrogen responsive genes. At longer exposures, these estrogen responsive genes initiate the transcription of a larger set of secondary transcription factors. These transcription factors then promote proliferation and suppress apoptotic genes.

We were able to observe the evolution of enrichment over increasing exposures by moving away from using traditional p-value and fold-change cutoffs to define lists to be used for calculating enrichment. These cutoffs do not account for low-level, diffuse patterns of gene expression that can characterize early time-points or low dose responses to exposure. Using the entire dataset instead of a limited set of highly expressed genes allowed us to investigate the cellular response at 2, 4 and 8 hours post-treatment, where the number of differentially expressed genes did not yield any enrichment information regarding either transcription factor binding or cellular processes. Additionally, IDEA allowed us to obtain results at conditions where array normalization and experimental noise would have severely decreased the utility of traditional enrichment methods. Because we are capturing information contained in relative expression of genes with respect to each other as opposed to some external cutoff, we will be better able to compare enrichment results across multiple experiments, which mitigate concerns about comparing functional enrichment results across multiple estrogen receptor studies in the NCBI Gene Expression Omnibus. Along with significantly enriched pathways and transcription factors, IDEA also provided us with the number of genes needed to achieve that enrichment (Supplementary Figure).This gives some insight into the strength of the signal in the data as it unfolds over time and dose, which can be useful for both experimental design and other bioinformatics approaches, which require dimensionality reduction.

In conclusion, IDEA provides us with a framework for observing patterns with gene expression studies and can provide a viable tool to investigate mode of action for multiple chemicals of the same class. The similarity of results with WGCNA is reassuring and these methods complement each other in the effort to provide a more nuanced characterization of estrogen’s PoT. The new approach lends itself for the initial identification of candidate PoT, which could then be followed by more targeted experiments on the path to a Human Toxome (Bouhifd et al., 2015) and a systems toxicology approach (Hartung et al., 2012).

## Supplemental Figures

**Supplementary Figure A.**
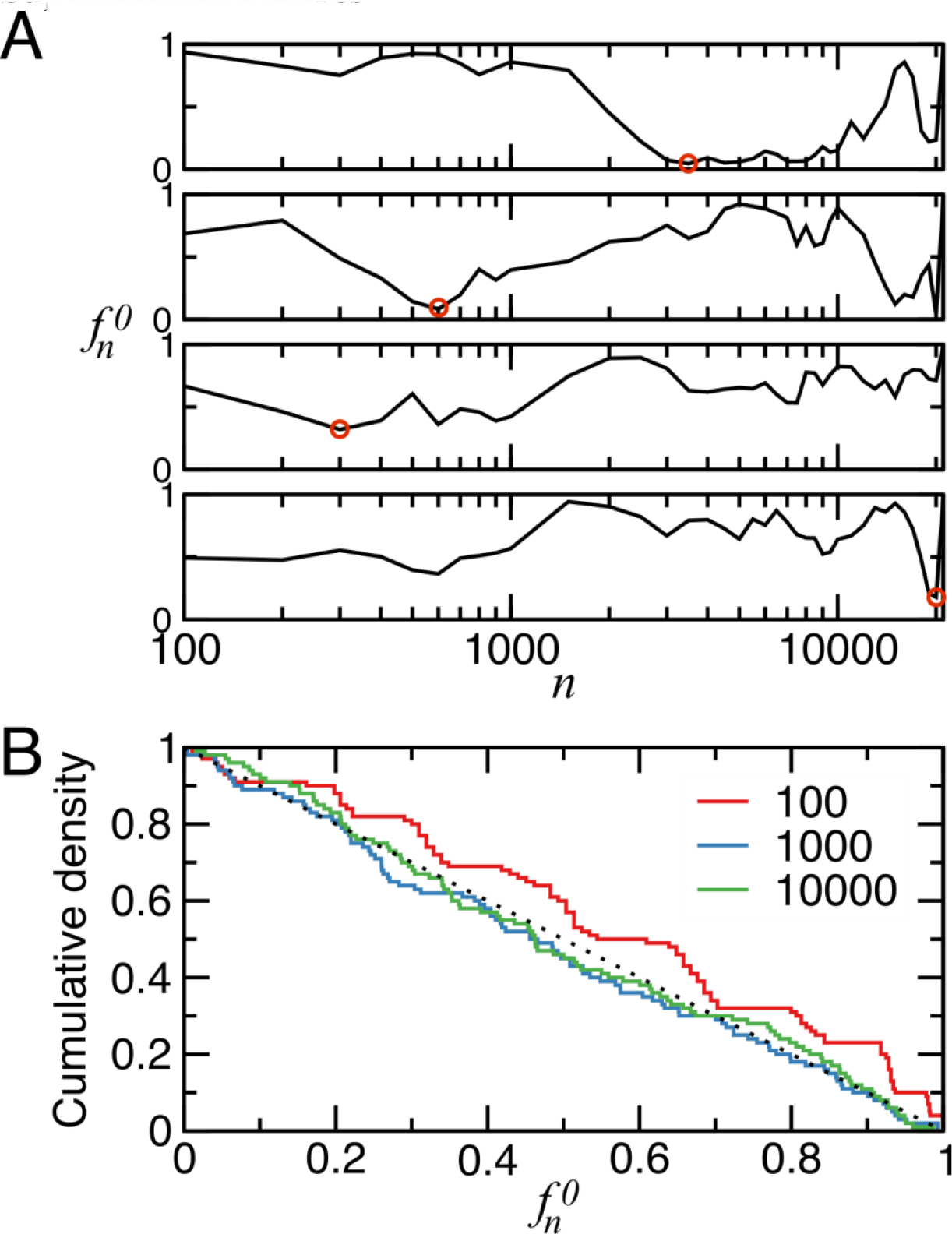
Characterization of the **IDEA** null model. (A) Four representative simulations of *f_n_^0^* for the association of up-regulated genes with ERα. Circles denote *t^0^* for each simulation. (B) Null model test statistics must be uniformly distributed in a valid Monte Carlo hypothesis test. Ensembles of *f_n_^0^* for «=100 (red), «=1000 (blue), and «=10,000 (green) are uniformly distributed (black dashed line) between 0 and 1.

**Supplementary Figure B.**
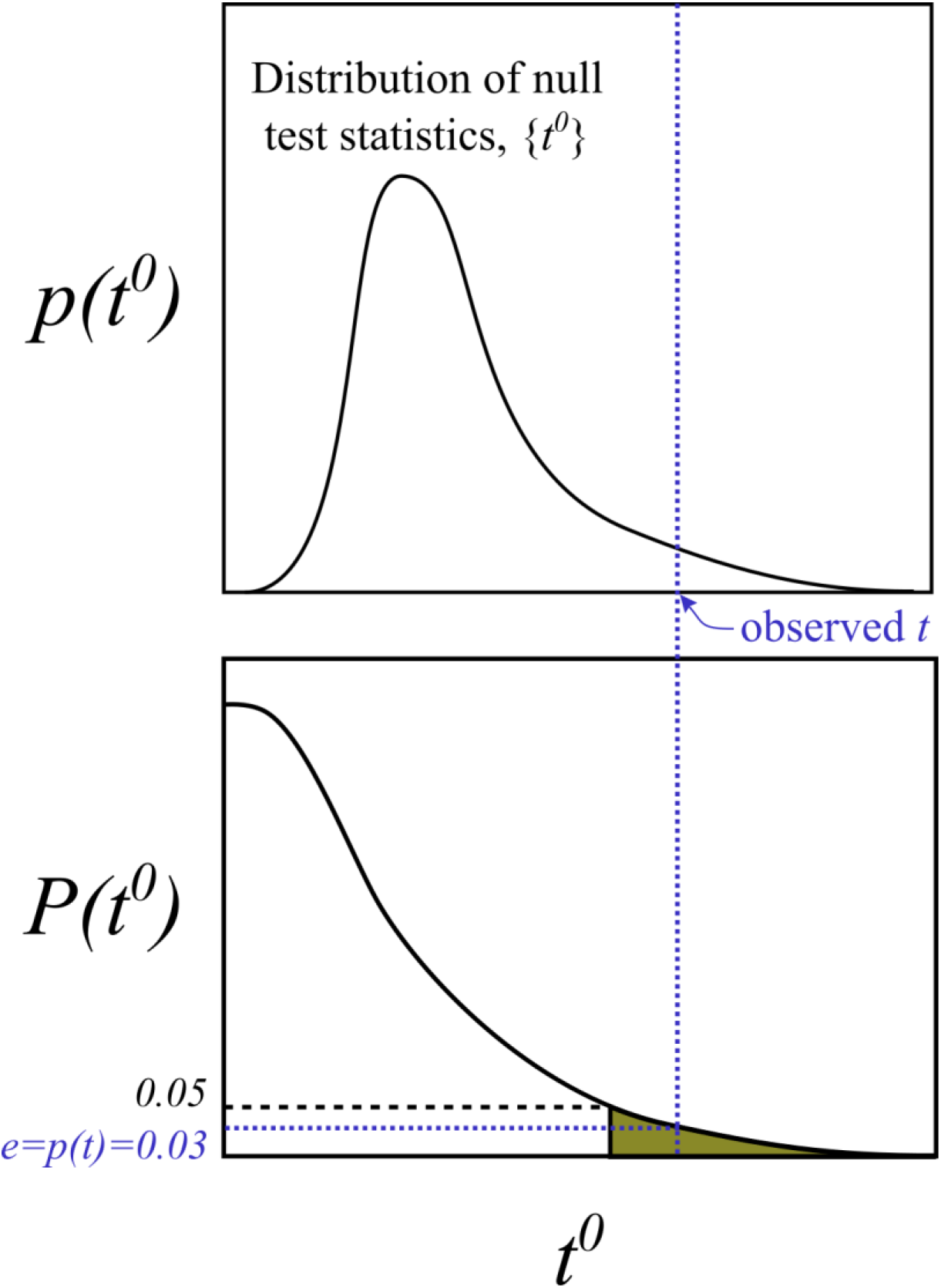
Successive simulations of *f_n_^0^* give rise to an ensemble of null model test statistics, {*t^0^*}. (A) Significance of enrichment, *e*, is based on the relationship between the observed test statistic *t* and {*t*^0^}. The probability of observing a test statistic more extreme than *t* is the cumulative density of {*t*^0^} evaluated at *t*.

**Supplemental Figure S3.**
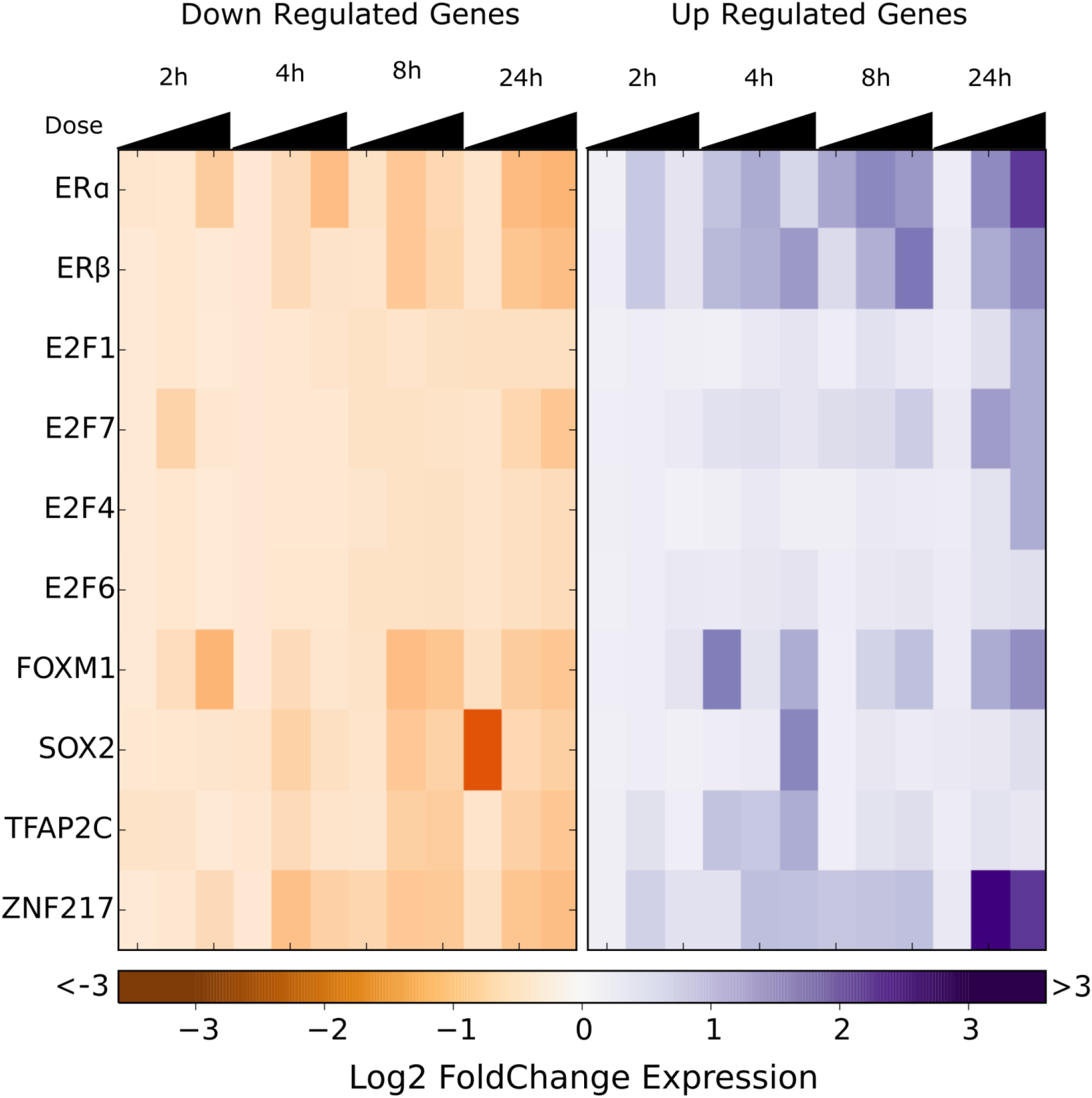
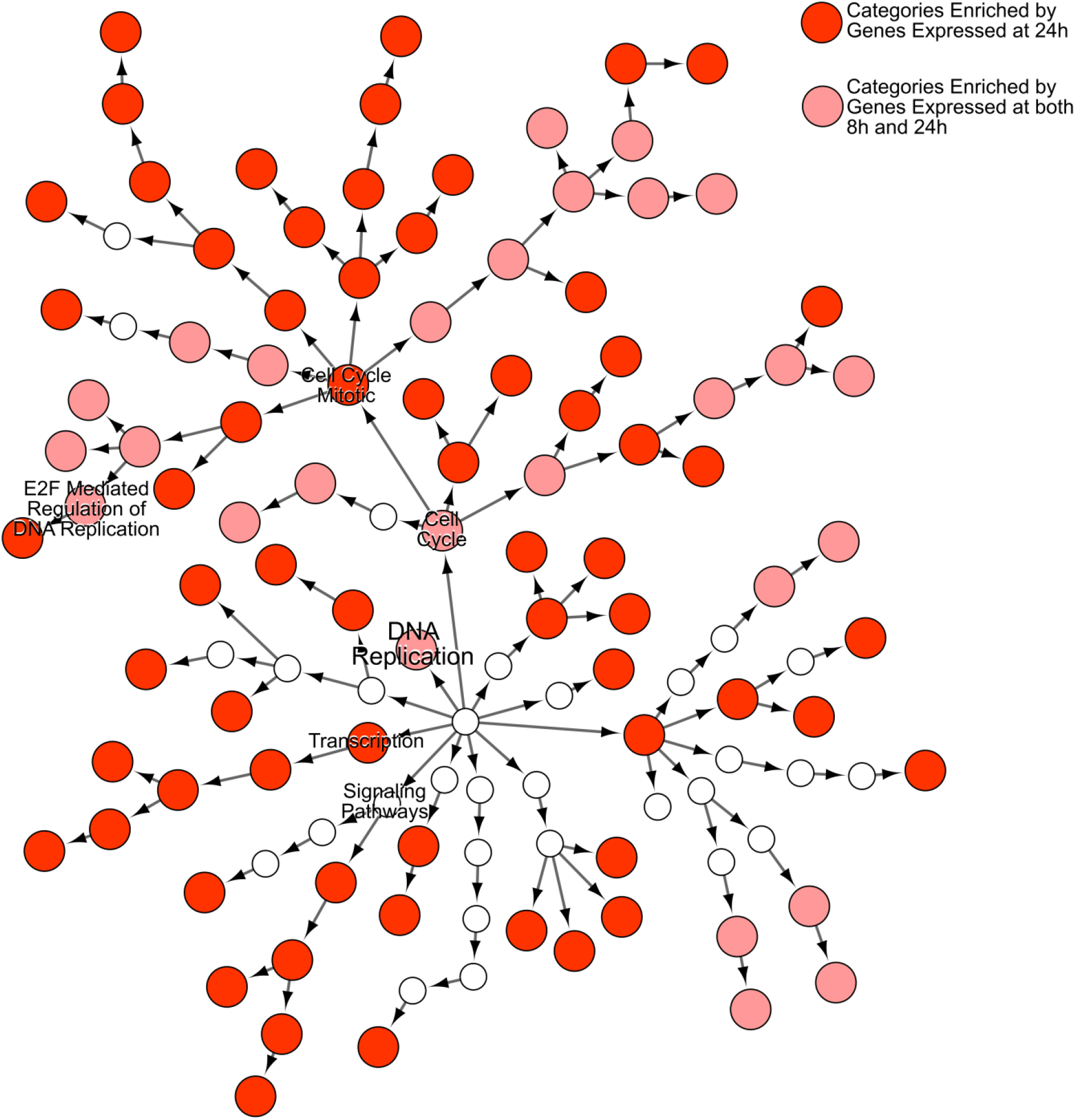
Reactome Enrichment using IDEA. Expression for genes regulated by key transcription factors. Significant enrichment of cell cycle related categories is observed only at 8 and 24h post exposure. A) Mean expression across all genes A) down-regulated and B) up-regulated by a given transcription factor and contributing to peak enrichment using IDEA.

**Supplementary Table T1:**
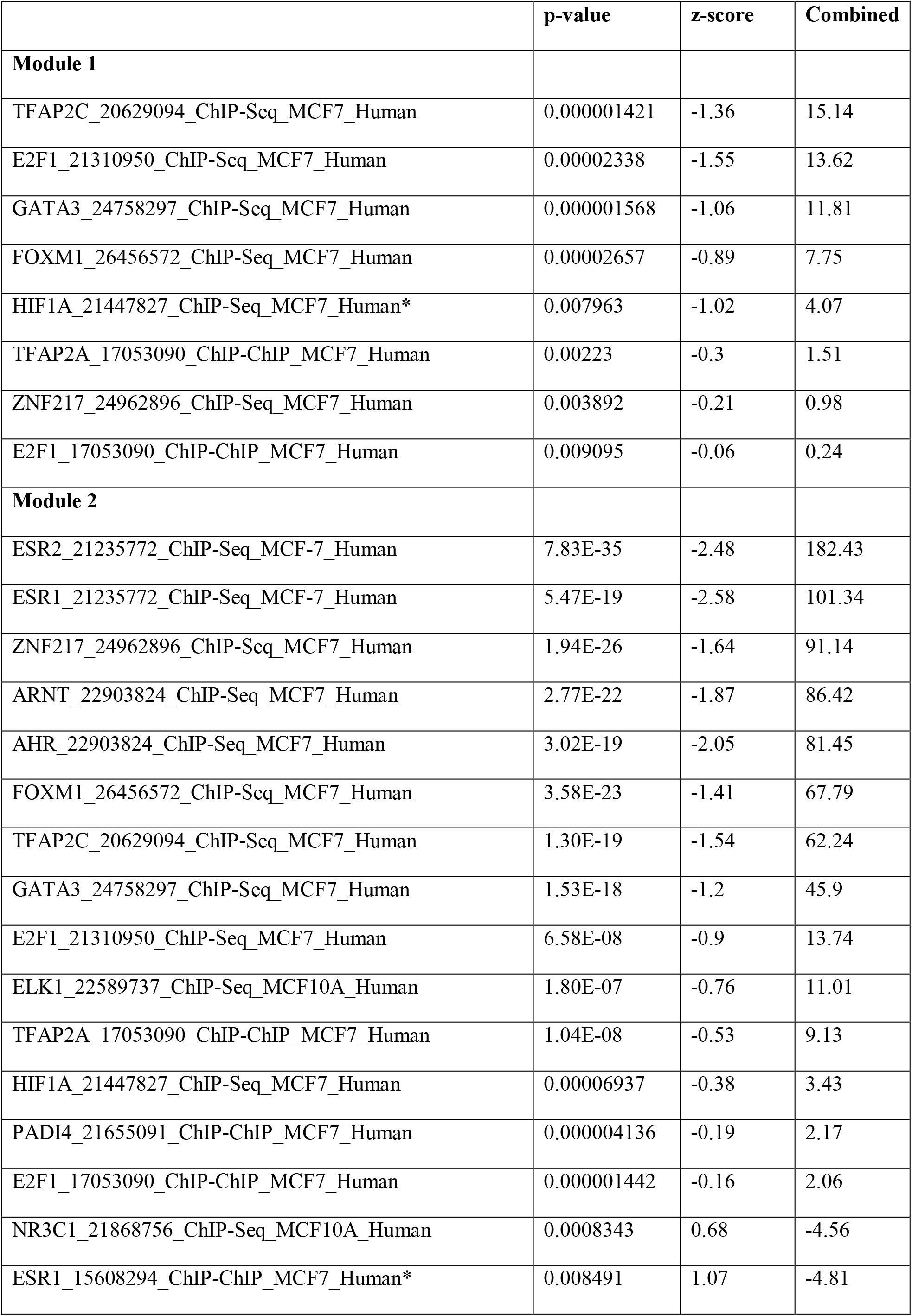
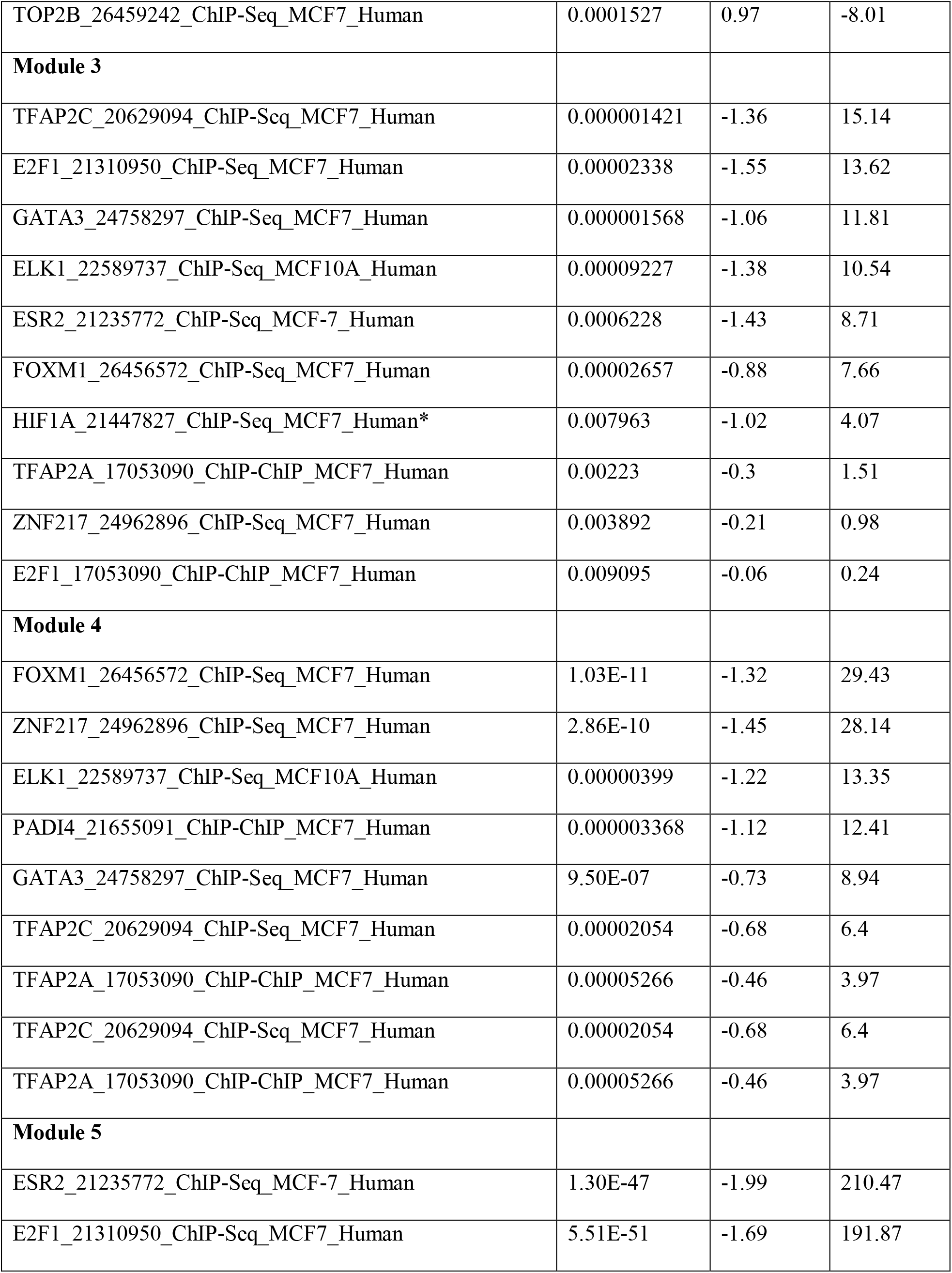
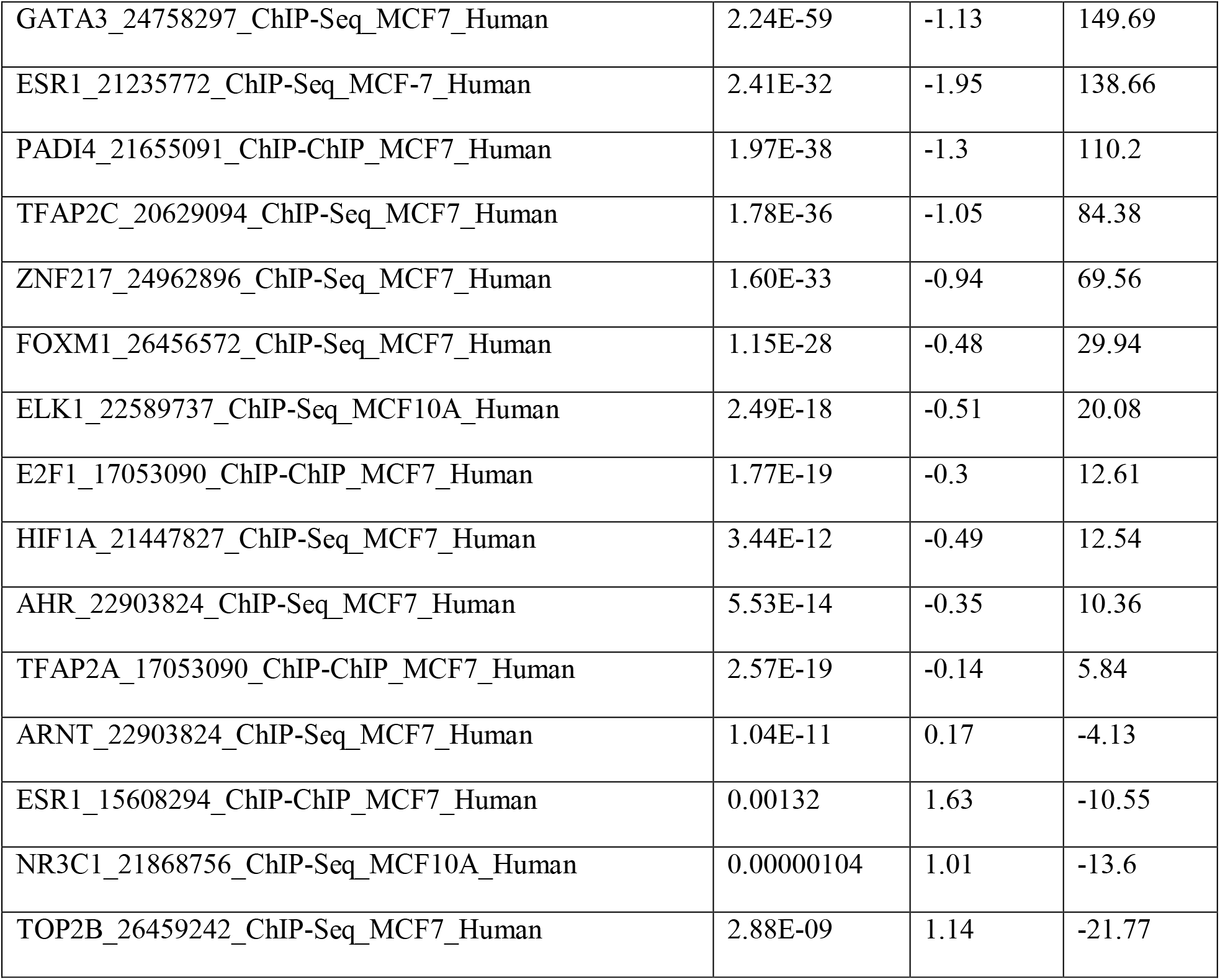
Transcription Factors Identified from Modules Derived from WGCNA of 8 hour dose-response curve.

## Footnotes

1 http://ntp.niehs.nih.gov/pubhealth/evalatm/test-method-evaluations/endocrine-disruptors/

